# Head down tilt 15° increases cerebral perfusion before recanalization in acute ischemic stroke. A pre-clinical MRI study

**DOI:** 10.1101/2024.08.02.606281

**Authors:** Simone Beretta, Davide Carone, Tae-Hee Cho, Martina Viganò, Susanna Diamanti, Jacopo Mariani, Francesco Andrea Pedrazzini, Elisa Bianchi, Cristiano Pini, Radu Bolbos, Marlene Wiart, Carlo Ferrarese, Fabien Chauveau

## Abstract

We investigated the therapeutic effect of head down positioning at −15° (head down tilt; HDT15) on cerebral collateral flow and infarct growth in a rat model of large vessel occlusion (LVO) stroke, using multi-modal MRI. Twenty-eight Wistar rats were randomly assigned to HDT15 or flat position for 60 minutes, starting 30 minutes after occlusion of the middle cerebral artery, followed by reperfusion. The perfusion shift analysis, comparing post- versus pre-treatment voxel-level changes in time-to-peak perfusion maps, showed a significant increase in cerebral perfusion in the HDT15 group (common odds ratio 1.50; 95% CI 1.41-1.60; p < 0.0001), but not in the flat group (common odds ratio 0.97; 95% CI 0.92-1.03; p = 0.3503). Infarct growth at 24 hours was + 31.4% in the flat group (343 versus 250 mm^3^; 95% CI 2.4 to 165.1; p = 0.0447) and + 15.4% in the HDT15 group (224 versus 192 mm^3^; 95% CI -26.9 to 85.9; p = 0.2272). Our findings indicate that HDT15 acutely increases cerebral perfusion in LVO acute ischemic stroke and provides a tissue-saving effect before recanalization. Further research is needed to develop HDT15 as an emergency therapy to acutely increase collateral flow in ischemic stroke prior to recanalization therapy.

## INTRODUCTION

The benefit of recanalization therapies in acute ischemic stroke is closely related to the subject-specific time window of the ischemic penumbra, whose evolution from salvageable tissue to irreversible infarction largely depends on the residual blood flow provided by cerebral collaterals.^1–3^

The degree of collateral blood flow predicts successful versus futile recanalization in patients treated with intravenous thrombolysis^4^ or mechanical thrombectomy.^5, 6^ Collaterals are an independent predictor of outcome even in untreated ischemic stroke patients with large vessel occlusion (LVO), providing residual blood flow to cortical ischemic areas despite persistent occlusion.^7, 8^

Investigating, developing and implementing a collateral therapeutic is a major objective in stroke research.^9^ The primary aim of enhancing cerebral collateral flow in acute ischemic stroke is to increase the amount of potentially salvageable penumbral tissue and expand the tissue time window, increasing the efficacy of recanalization therapies. The ideal collateral therapeutic should not be only effective, but also easy to apply, rapidly active and safe.

Experimental studies from independent laboratories showed that head down tilt (HDT) ranging from −15° to −60° improved stroke outcome in rodent stroke models and increased cerebral blood flow measured by laser Doppler and two-photon microscopy.^10–13^

Clinical studies with head positioning has been mostly focused on lying flat (0° positioning), compared to +30°.^14^ Despite increased cerebral blood flow velocity measured by transcranial Doppler, near-infrared or diffuse correlation spectroscopy has been consistently associated with a lower head position,^15–21^ a neutral effect on functional recovery was clearly shown in the HeadPoST trial.^22^

No previous study explored the effect of HDT in acute ischemic stroke using neuroimaging, either in animals or in patients. In the present study, we performed a preclinical randomized trial to investigate the acute effect of HDT at −15° (HDT15) on MRI-based cerebral perfusion metrics and early infarct growth in a rat stroke model of LVO.

## MATERIAL AND METHODS

### Experimental design and sample size determination

The protocol of this study was pre-registered at preclinicaltrials.eu, identifier PCTE0000198. SB had full access to all the data in the study and takes responsibility for its integrity and the data analysis. Data and neuroimages regarding the primary outcome measure have been made publicly available at the OpenNeuro repository and can be accessed at [https://doi.org/10.18112/openneuro.ds004962.v1.0.0]. Further data is available from the corresponding author upon reasonable request. The experimental protocol was approved by the review board "Comité d’éthique pour l’Expérimentation Animale Neurosciences Lyon" (CELYNE - CNREEA number: C2EA – 42), and by the Committee on Animal Care of the University of Milano-Bicocca, in accordance with the national guidelines on the use of laboratory animals (Decree 2013-118 in France; D.L. 26/2014 in Italy) and the European Union Directive for animal experiments (2010/63/EU). Authorization for the use of animals for this project was obtained under project license from the French Ministry of Higher Education and Research (15529- 2018061512184831v2 and 32924-2021091015062327v4), and the Italian Ministry of Health 1056/2020-PR. Procedures for the use of animals were reported in accordance with guidelines from the National Institutes of Health and Animal Research Reporting of In Vivo Experiments (ARRIVE).

In a randomized, controlled, parallel group, pre/post interventional trial, consecutive animals undergoing successful middle cerebral artery (MCA) occlusion was used to explore the effect of HDT15 on cerebral perfusion in the ischemic hemisphere, compared to flat position. The primary outcome was change in cerebral perfusion, assessed by relative time-to-peak (TTP), after 1 hour of treatment. The secondary outcome was infarct growth in the first 24 hours.

Additional perfusion metrics (cerebral blood flow [CBF] and cerebral blood volume [CBV]) were considered exploratory outcomes and included as supplementary results.

Infarct volume, functional neurological outcome and stroke-related mortality at 24 hours were considered exploratory outcomes and included as supplementary results, since they have already been published in a pooled analysis with other two experimental series.^11^

Our previous data indicated that cerebral perfusion slightly decreases during a 90-minutes MCA occlusion in rats maintained in the flat position (TTP change +2%), with a standard deviation of 35%. An increase in cerebral perfusion corresponding to a TTP change −30%, due to treatment, was considered to be a response of substantive translational significance. Under these assumptions, we planned a study of a continuous variable (TTP) from matched pairs (pre- and post-treatment; two groups, HDT15 and flat) with a sample size of 28 animals (14 matched pairs per treatment group), with an expected power of 0.80 and type I error of 0.05 to refuse the null hypothesis that the response difference is zero (two-sided significance level).

### Randomization, allocation concealment and blinding

A simple unrestricted randomization was performed using an online random number generator (www.random.org). In order to reach the calculated sample size, consecutive rats were randomized until each group (HDT15 or flat position) reached the minimum number of 14 animals. Treatment group was disclosed after successful MCA occlusion to guarantee allocation concealment to the experimental surgeon. Outcome assessment, including image analysis and functional neuroscore, was performed by investigators blinded to treatment.

### Animals and surgery

Adult male Wistar rats (ILAR code Crl:WI (Han); weight 298 ± 14 g; total n = 40; Charles River, L’Arbresle, France) were anesthetized with 3% isoflurane in O2 /N2O (1:3), and maintained with 1.5% isoflurane. A catheter was placed on one vein of the tail. Occlusion of the origin of the right MCA was induced transiently for 90 minutes with a reperfusion period of 24 hours. Briefly, a silicone-coated filament (diameter 0.39±0.02 mm, Doccol Corporation, Redlands, CA, USA), was introduced in the right external carotid artery and pushed through the right internal carotid artery to occlude the origin of the right MCA.

Common carotid artery was permanently occluded by ligation immediately before the insertion of the filament. Proximal branches of the external carotid artery (occipital artery and cranial thyroid artery) were electrocoagulated during the surgical procedure, before the insertion of the filament. Immediately after MCA occlusion, rats were transferred in the MRI scanner for the image acquisition protocol and treatment (see below), under isoflurane anesthesia. During surgery and MRI acquisition, the core temperature of 37°C was controlled by a rectal thermometer connected to a feedback- controlled heating pad. Respiration rate was controlled visually during surgery and through a respiratory sensor placed on the abdomen during MRI. Reperfusion was obtained by gentle removal of the endovascular filament, followed by electrocoagulation of the external carotid artery stump and cervical wound closure. After reperfusion, rats were allowed to recover from anesthesia and had free access to food and water. After 24 hours from the onset of ischemia, animals were assessed for neurobehavioral score, then they were anesthetized with isoflurane for the follow-up MRI session. After completing the last MRI sequence, animals were euthanized by intracardiac injection of Pentobarbital (Euthasol), under isoflurane anesthesia.

A detailed description of the housing, perioperative care and mortality is available in the Supplemental Material.

### Inclusion and exclusion criteria

Six rats were excluded from the experimental series for early death due to peri- procedural subarachnoid hemorrhage (SAH) occurred during MCA occlusion. SAH was suspected by MRI appearance on the pre-treatment T2 sequence (hypointense lining of peri-cisternal spaces) and verified by necropsy (macroscopic bleeding around the circle of Willis). Three rats were excluded for unsuccessful MCA occlusion, verified by MR angiography (persistence of symmetric signal in the right and left MCA). Two rats were excluded for anesthesia-related death before completing the perfusion MRI protocol.

One rat was excluded for combined MCA + anterior cerebral artery occlusion, verified by angio-MRI and DWI sequences (massive hemispheric infarction, exceeding MCA territory).

All rats which were subjected to successful MCA occlusion (verified by loss of MCA signal on MR angiography and presence of hyperacute ischemic lesion on DWI sequences) and completed the perfusion MRI protocol (n=28) were included for the primary outcome analysis.

### Application of HDT15

HDT15 was applied for 60 minutes, starting 30 minutes after MCA occlusion, to the rats lying in the prone position on the MRI scanning bed, using a custom-made, 15° tilted nylon platform. During HDT application, the rat’s head remained still in the flat position, while the rest of the body was tilted, to prevent image distortion in brain MRI acquisition and allow a direct comparison between pre- and post-treatment. After 60 minutes, rats were returned to the flat position and MCA reperfusion was obtained.

### Image acquisition and processing

Immediately after occlusion, the rats were placed in a dedicated plastic bed equipped with a stereotactic holder (Bruker Biospec Animal Handling Systems, Germany) and maintained under gaseous anesthesia delivered via a cone mask throughout the MRI protocol.

MRI acquisitions were performed on a horizontal 7T Bruker Biospec MRI system (Bruker Biospin MRI GmbH, Germany) with a set of 400 mT/m gradients, controlled by a Bruker ParaVision 5.1 workstation. A Bruker birdcage volume coil (outer diameter = 112 mm and inner diameter = 72 mm) was used for signal transmission and a Bruker single-loop surface coil (25 mm diameter), positioned on the head of the animal to target the brain, for signal reception.

For each rat, the following MRI protocol was performed (Figure 1 for general outline; Table 1 for timings; Table S1 for additional details):

- a first dynamic susceptibility contrast-enhanced perfusion weighted imaging (DSC-PWI pre), during MCA occlusion, before treatment
- baseline anatomical T2 weighted imaging (T2 baseline)
- apparent diffusion coefficient (ADC) map (early infarct), approximately 45 minutes after MCA occlusion
- MR angiography, during MCA occlusion
- a second dynamic susceptibility contrast-enhanced perfusion weighted imaging (DSC-PWI post), during MCA occlusion, approximately 60 minutes after treatment, immediately before reperfusion
- a follow-up T2 weighted imaging (T2 final), 24 hours after reperfusion

**Figure 1.**
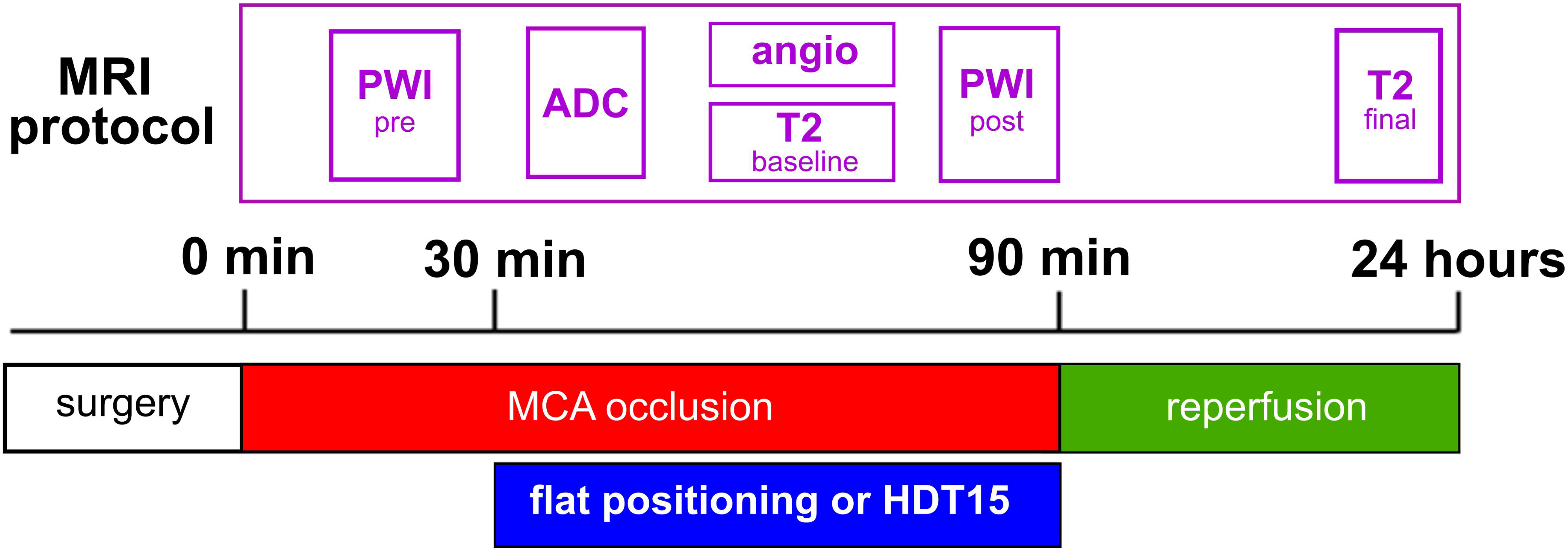
Experimental outline. Schematic representation of the experimental procedure, treatment application and MRI protocol.

**Table 1.**
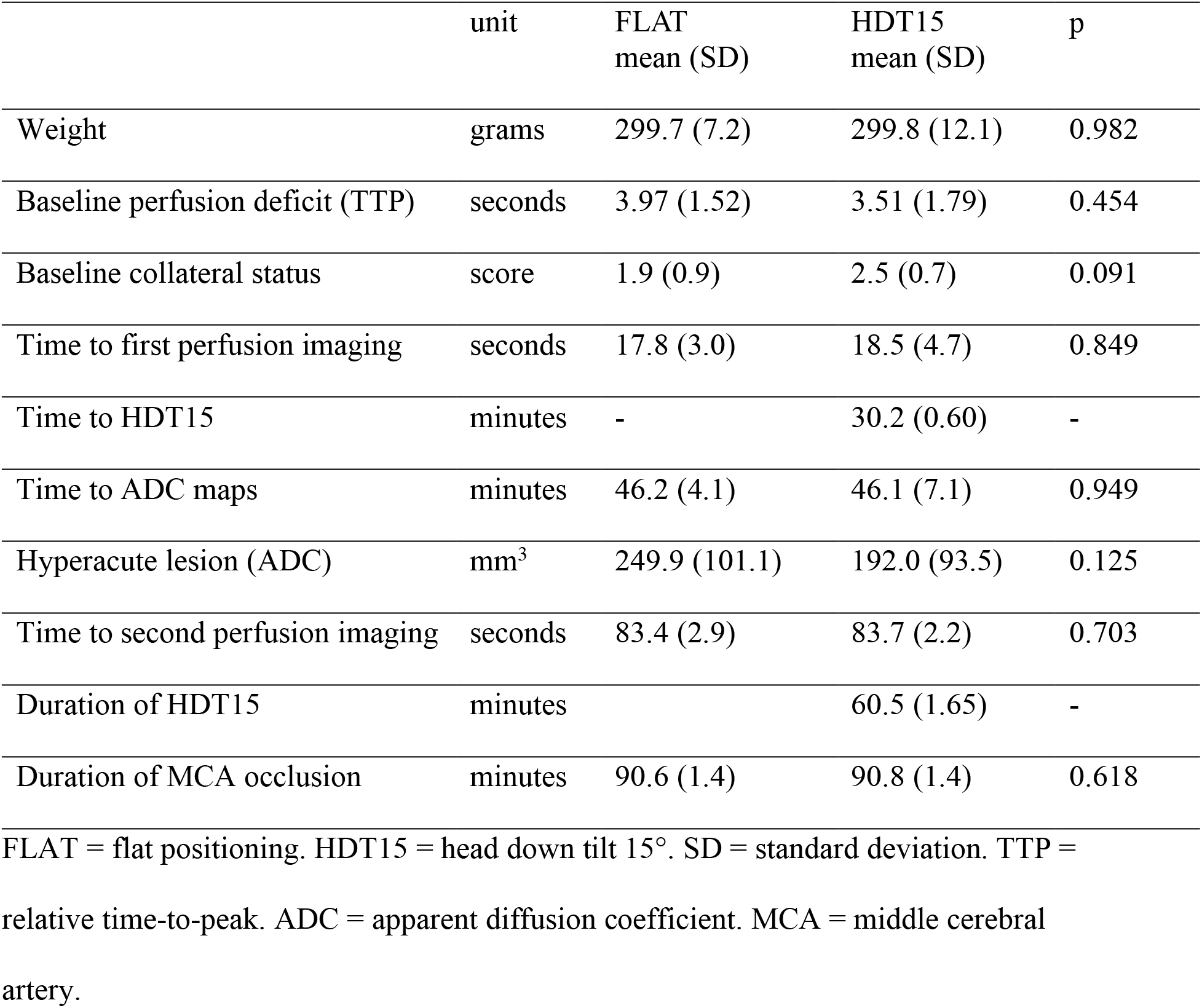
Baseline characteristics of the study groups.

All image analyses were performed using the Oxford Centre for Functional MRI of the Brain (FMRIB) software library (FSL; https://fsl.fmrib.ox.ac.uk/fsl/fslwiki). A detailed description of image processing, region of interest (ROI) and masking in available in the Supplemental Methods.

### Perfusion analysis

TTP maps were generated off-line using an in-house Matlab code. CBF and CBV maps were generated using VERBENA^23^ (Singular Value Decomposition deconvolution option). The arterial input function time course was extracted from the AIF mask.

All perfusion maps were expressed relative to the contralateral hemisphere (TTP maps were subtracted for the median TTP values of the contralateral hemisphere, whilst CBF and CBV maps were normalized by dividing for the median value of the contralateral hemisphere).

Perfusion values were then extracted at a voxel level to enable composite voxel-wise analysis across animals and to estimate the voxel-wise mean in the ischemic hemisphere.

### Assessment of baseline collateral score

Baseline collateral status was grade from 0 (no collaterals) to 4 (excellent collaterals) by an experienced vascular neurologist using the American Society of Interventional and Therapeutic Neuroradiology/Society of Interventional Radiology (ASITN/SIR) collateral score^24, 25^ on DSC-PWI sequences.

### Neurologic outcome assessment

Garcia neuroscore was used to assess global neurological functioning at 24 hours on an ordinal scale from 3 (most severe) to 18 (no deficit) and required scoring spontaneous movement, sensory function and motor function.^26^ Stroke-related mortality was assessed at 24 hours and was expressed as 0 for the Garcia neuroscore and separately as a dichotomous variable.

### Statistical analysis

Data were analyzed using SAS 9.4 (SAS Institute, Cary, NC, USA) and GraphPad Prism 9.1.1 (GraphPad Software, Boston, MA, USA). Values were expressed as mean +/- standard deviation (SD). Relative percent difference was calculated as the difference between two values, divided by the average of the two values, shown as a percentage.

The two-group analysis on baseline characteristics was done by means of Mann-Whitney test.

Changes in perfusion were explored in three different ways: *ROI analysis:* for each treatment-group, changes in perfusion were explored by comparing the mean TTP in the entire ischemic hemisphere across time, for each animal, by means of paired t-test. This allowed to estimate an overall treatment effect and determine if changes were not driven by one specific animal.

*Composite group level voxel-wise analysis:* all voxels in the ischemic hemisphere were combined and compared across time and treatment groups. This approach allowed to maximize statistical power. Voxel-level perfusion analyses (TTP) were performed using a multilevel linear mixed model with TTP as dependent variable, treatment, scan (pre- and post-treatment) and treatment*scan interaction as independent variables and an unstructured variance covariance matrix. The model was also adjusted for the baseline collateral score. Mean TTP value was estimated in HDT and flat group pre- and post- treatment, along with differences between post- and pre-treatment (delta). Treatment effect was quantified as the difference between post and pre-treatment values between the two treatment groups, which is tested by the treatment*scan interaction term.

Perfusion shift analysis (TTP) was performed using a multilevel multinomial logistic regression model in the HDT and flat group separately, comparing post to pre- treatment. TTP was categorized (<2, 2-4, 4-6, 6-8, 8-10, 10-12, 12-14, >14) and included in the model as dependent variable. Independent variables were scan and collateral score as adjustment covariate. An adjusted common odds ratio, with 95% CI was estimated and reported for each treatment group.

*Voxel-wise spatial analysis:* to localize significant changes in cerebral perfusion over time at a group level, a voxel-wise two-sample paired t-test was performed on the TTP perfusion maps using a general linear model. For non-parametric permutation testing, FSL’s randomise (version 2.9) was used with 5000 permutations. Statistical thresholding was performed with FSL’s threshold-free cluster enhancement (TFCE) and a family-wise error rate (FWE) of p smaller than 0.05. This analysis allowed to explore the presence of common spatial patterns in perfusion change within the two treatment groups.

Infarct growth was analyzed for each treatment-group by comparing the volume of the ischemic lesion on hyperacute ADC sequence (at 46 minutes; see Table 1) with the one on T2 sequence at 24 hours, for each animal, by means of paired t-test.

The two-group analysis on mortality was done by means of Fisher’s exact test. The two- group analysis on neuroscore and infarct size was done by means of unpaired t-test. A p-value of less than 0.05 was considered significant.

## RESULTS

### Study population

A total of 28 consecutive rats were randomized and included in the analysis, 14 rats were allocated to flat positioning and 14 rats were allocated to HDT15. The main baseline characteristics and timings showed no significant difference between the two groups (Table 1). HDT15 group displayed a tendency towards better baseline collateral status, compared to the flat group (2.5 +/- 0.7 versus 1.9 +/- 0.9; p = 0.091). ADC maps were acquired at approximately 46 minutes after MCA occlusion and showed a high variability in both groups. HDT15 was applied at 30.2 +/- 0.6 minutes after MCA occlusion and maintained for 60.5+/-1.6 minutes until reperfusion.

### Acute effect of HDT15 on cerebral hemodynamic during MCA occlusion

The ROI-level analysis showed that rats in the flat group had a mild, non-significant improvement of cerebral perfusion over time (TTP absolute difference −0.24 sec; 95% CI −0.73 to 0.23; relative percent difference 6.4%; p = 0.2884; Figure 2A). Conversely, application of HDT15 significantly improved cerebral perfusion, compared to pre- treatment (TTP absolute difference −0.86 sec; 95% CI −1.56 to −0.17; relative percent difference 28.2%; p = 0.0182; Figure 2B).

**Figure 2.**
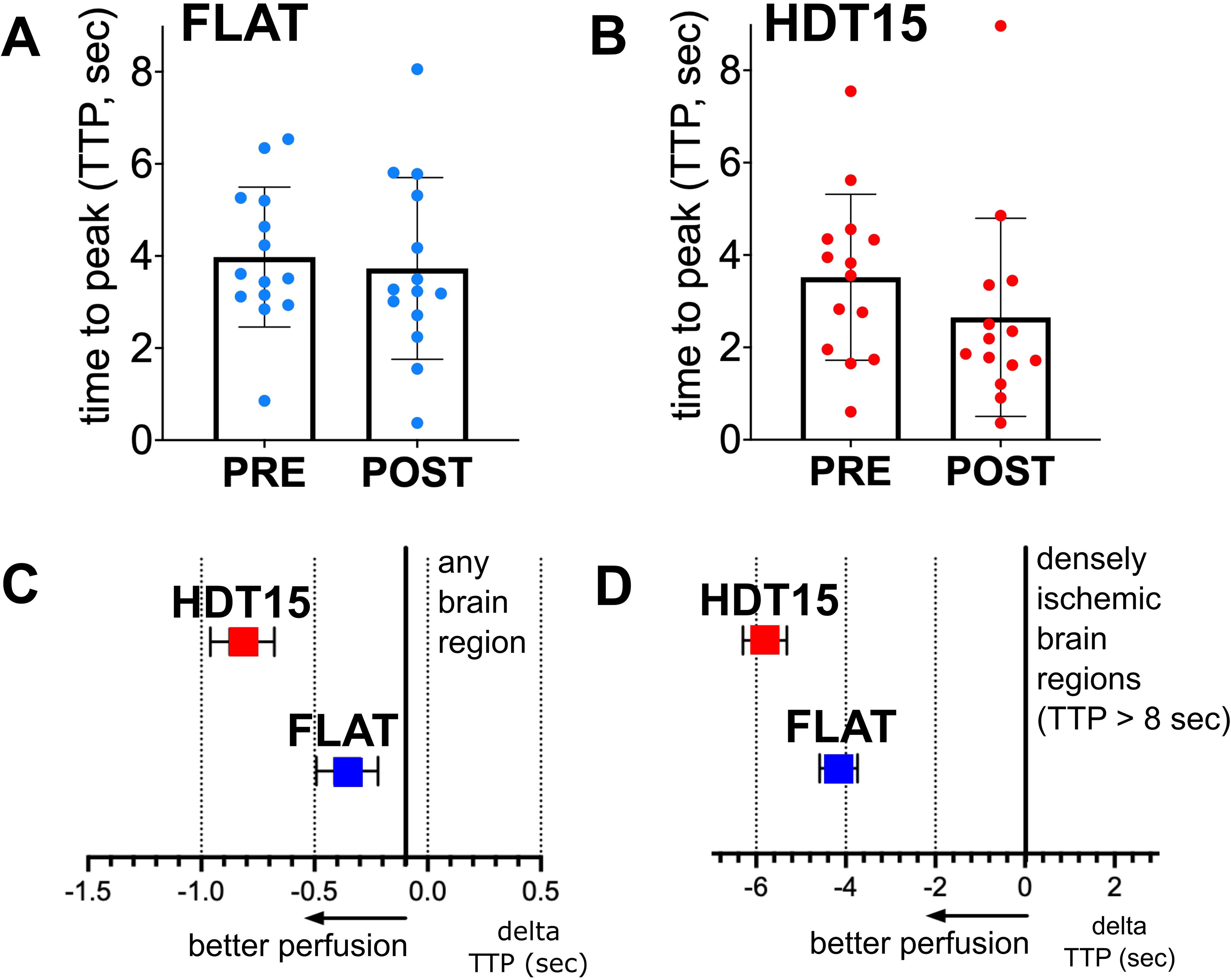
Quantitative changes of cerebral perfusion during MCA occlusion, before and after HDT15 application. TTP values during MCA occlusion were averaged across the entire ischemic hemisphere (ROI-level), comparing before and after flat positioning (**A**) and HDT15 (**B**) for 1 hour. Voxel-level quantification of the effect of flat positioning and HDT15 on cerebral perfusion during MCA occlusion, assessed as the difference (delta) of TTP over time, is shown for any brain region (**C**) or densely ischemic brain regions (TTP > 8 seconds; **D**) in the ischemic hemisphere.

The voxel-level analysis, comparing post- versus pre-treatment for each group, adjusted for the baseline collateral score, showed that cerebral perfusion increased in the HDT15 group, compared to the flat group, across the entire ischemic hemisphere (-0.81 versus - 0.35 sec; absolute difference 0.46 sec; 95% CI −0.65 to −0.26; relative percent difference 14.6% versus 6.3%; HDT15 versus flat, respectively; p < 0.0001; Figure 2C). The change in cerebral perfusion, in favor of HDT15, became larger when this analysis was focused the more densely ischemic areas (rTTP > 8 sec; −5.81 versus −4.16 sec; absolute difference −1.65 sec; 95% CI –2.29 to −0.99; relative percent difference 31.6% versus 22.3%; HDT15 versus flat, respectively; p < 0.0001; Figure 2D).

The voxel-wise analysis (Figure 3) showed a significant increase in cerebral perfusion over time in both HDT15 and flat groups, which co-localized in the same cortical and subcortical areas within the MCA territory in the ischemic hemisphere. However, the regional distribution of this hemodynamic effect was wider and displayed a greater magnitude in the HDT15 group.

**Figure 3.**
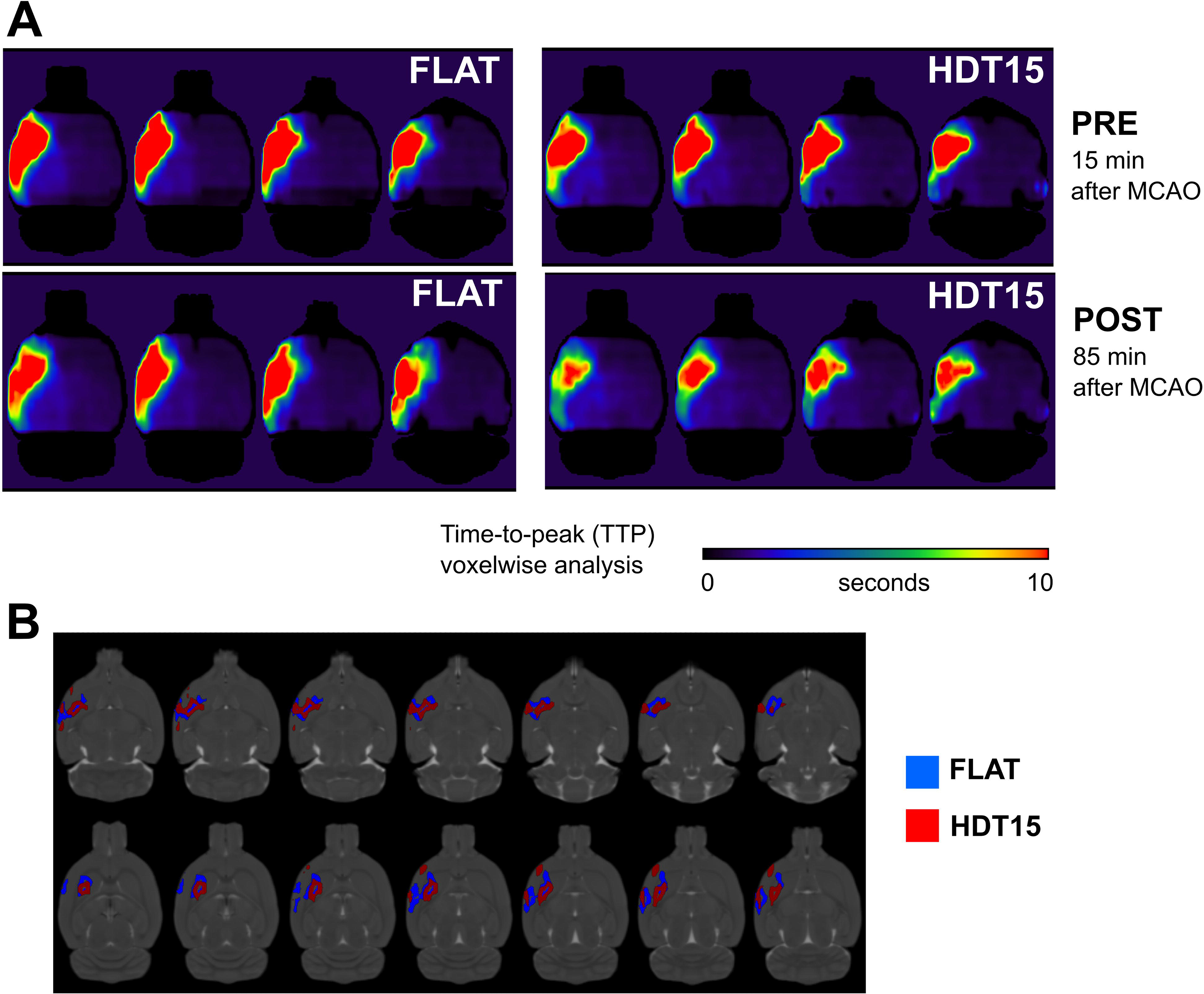
Changes in TTP (Voxelwise analysis) during MCA occlusion, before and after HDT15 application. (A) Spatial maps showcasing the average values of TTP for each treatment group (flat positioning: left; HDT15: right), before and after HDT. **(B)** Spatial maps showcasing areas where significant increase in perfusion (reduction in TTP) at a group level (blue: flat, red: HDT). Perfusion increased in the more peripheral regions of the perfusion deficit, suggesting recruitment of already existing collaterals.

The perfusion shift analysis (Figure 4), comparing post- versus pre-treatment for each group, adjusted for the baseline collateral score, showed a higher probability to shift towards lower TTP values (i.e. improve cerebral perfusion) in the HDT15 group (common odds ratio 1.50; 95% CI 1.41-1.60; p < 0.0001), whereas no significant shift in perfusion was observed in the flat group (common odds ratio 0.97; 95% CI 0.92- 1.03; p = 0.3503).

**Figure 4.**
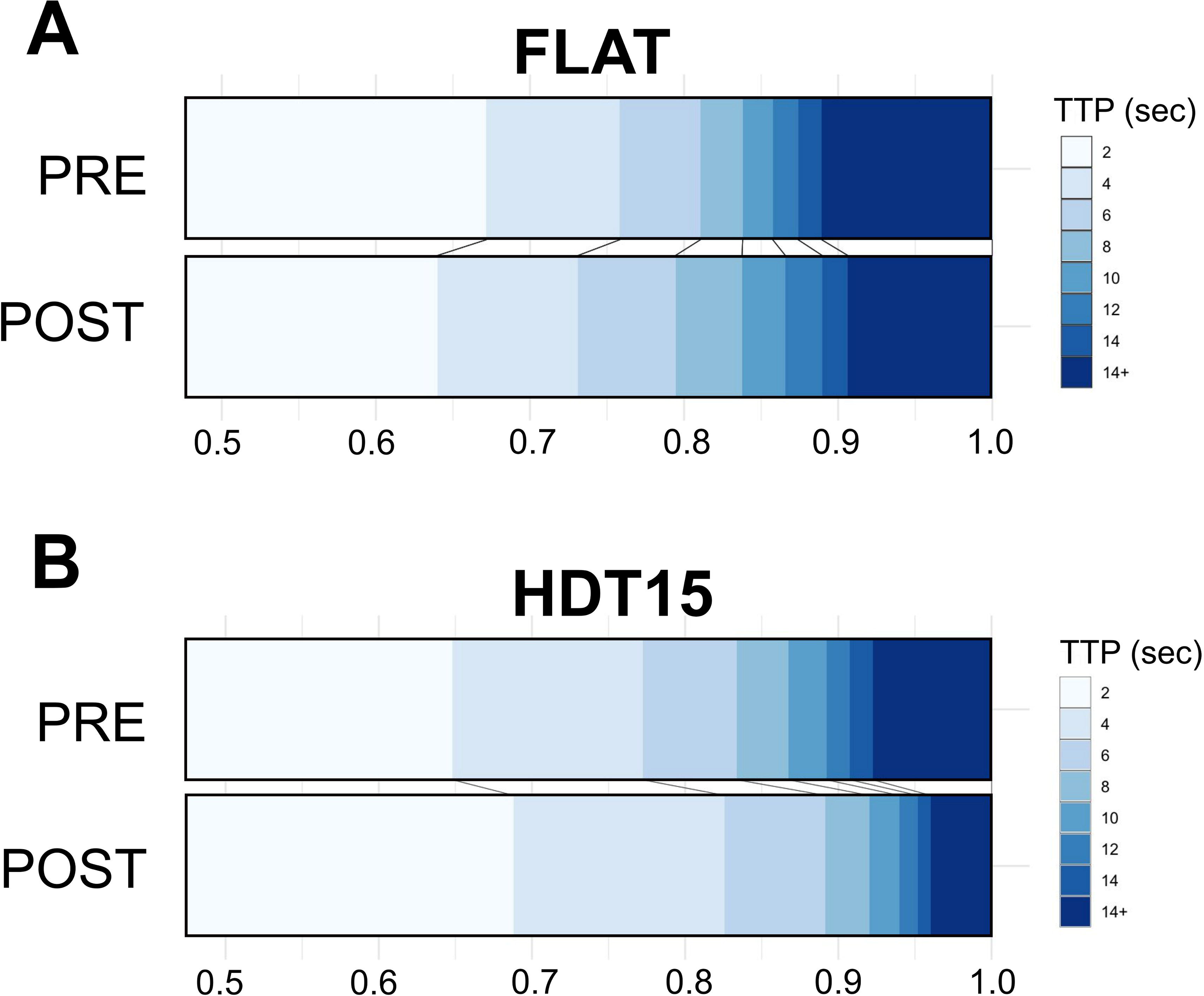
Perfusion shift analysis during MCA occlusion associated with HDT15. Voxel-level TTP values during MCA occlusion were categorized into mildly ischemic (< 2 sec), severely ischemic (> 14 sec) and six intermediate categories (every 2 sec). The distribution of TTP categories is shown, comparing before and after flat positioning (**A**) and HDT15 (**B**) for 1 hour, and it was considered an indicator of the perfusion shift associated with treatment.

The voxelwise analysis of CBF (Figure S1) showed a significant increase over time in the HDT15 group within the ischemic MCA territory, while no significant changes was observed in the flat group.

The voxelwise analysis of CBV (supplementary Figure 1) showed no significant change over time in either groups.

### Effect of HDT15 on infarct growth and stroke outcome

Infarct growth after MCA occlusion, comparing the hyperacute ADC lesion with the T2 lesion at 24 hours, showed a 31.4% growth in the flat group (relative percent difference; 343 versus 250 mm^3^; absolute difference 93 mm^3^; 95% CI 2.4 to 165.1; p = 0.0447) and a 15.4% in the HDT15 group (relative percent difference; 224 versus 192 mm^3^; absolute difference 32 mm^3^; 95% CI -26.9 to 85.9; p = 0.2272; Figure 5).

**Figure 5.**
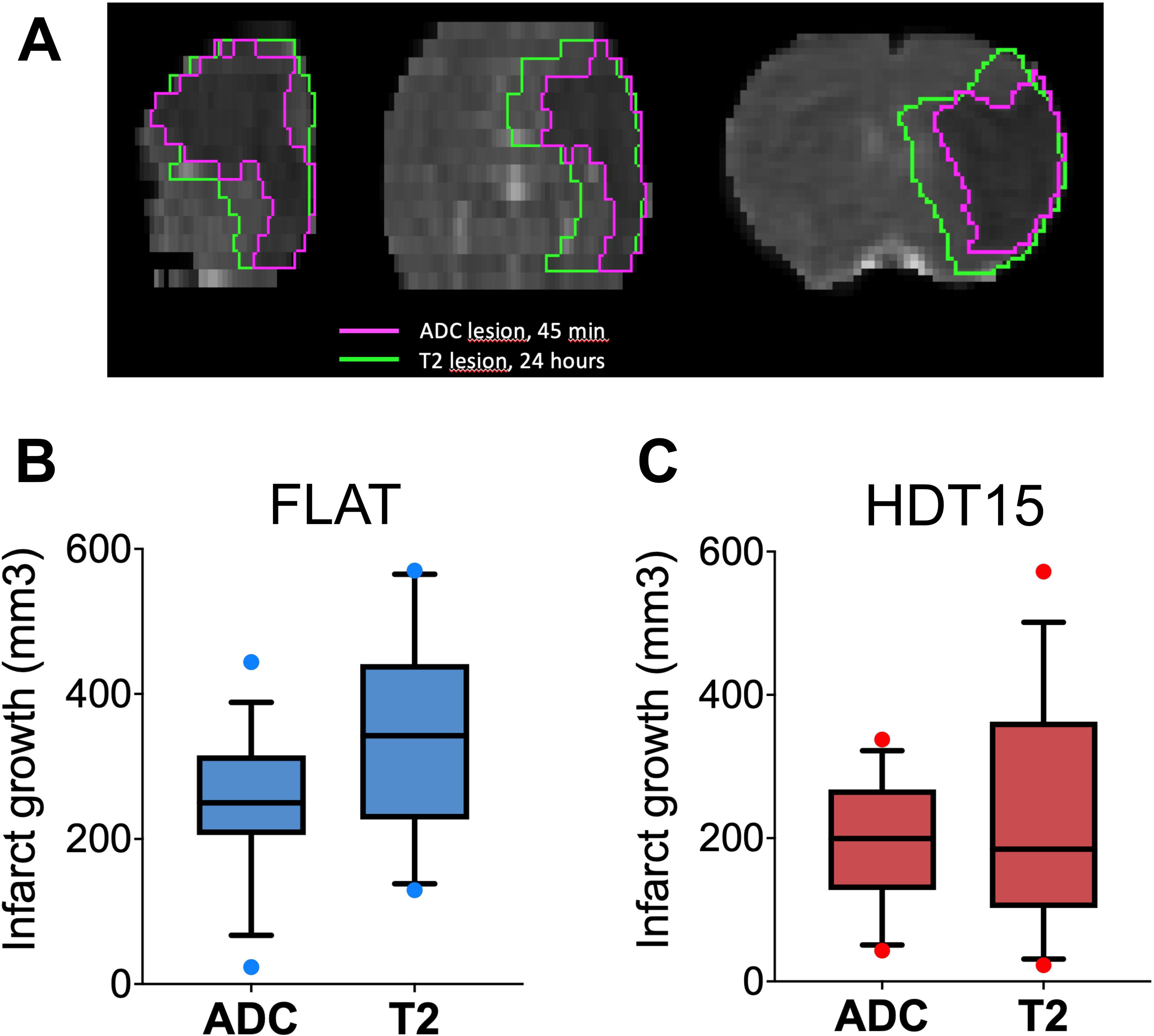
Infarct growth in the first 24 hours after transient MCA occlusion. Representative MRI images of a rat subjected to MCA occlusion, showing infarct growth (**A**) as the difference between the hyperacute ischemic lesion (ADC-positive, 45 min after occlusion; purple line) and the respective final infarct on T2-weighted images at 24 hours (green line). Infarct growth over time is shown for rats treated with flat positioning (**B**) and HDT15 (**C**).

Data on stroke-related mortality, neuroscore and infarct size at 24 hours are reported in Table S2. All stroke outcome measures showed a tendency in favor of HDT, compared to flat positioning (stroke-related mortality 21.4% versus 0%, p = 0.2222; Garcia neuroscore mean 7.1 versus 10.3, p = 0.0705; infarct volume mean 341 mm^3^ versus 224 mm^3^, p = 0.0575; flat versus HDT15, respectively).

## DISCUSSION

An evidence-based collateral therapeutic for acute ischemic stroke is urgently needed at a global level, to be applied before and maximize efficacy of intravenous thrombolysis and mechanical thrombectomy in the extended time window.^5,27,28^

The optimal timing for the application of a collateral therapeutic is the early phase of ischemic stroke, even in the pre-hospital setting, to enhance cerebral collateral blood flow prior to recanalization therapies.^29^ For its simplicity, low cost and feasibility, HDT15 represents a good candidate as a collateral therapeutic to be applied as soon as an acute ischemic stroke is suspected, even a few minutes after arterial occlusion.

Our preclinical randomized trial investigated the acute cerebral hemodynamic effect of HDT15 during MCA occlusion in rats, using perfusion imaging. Our findings indicate that HDT15 application produced a rapid improvement of cerebral perfusion measured by TTP, both at single rat level (ROI-based approach) and at group level (voxel-wise approach and voxel-level perfusion shift analysis). Interestingly, a modest increase in perfusion in the same cortical and subcortical areas was also observed in rats maintained in the flat position, although this effect was significantly smaller compared to rats treated with HDT15. This finding suggests that the hemodynamic effect of HDT15 application during MCA occlusion may act as a booster of the naturally occurring recruitment of collaterals in the ischemic brain areas.

Overall, the effect size of HDT15 positioning could be estimated approximately as a 15- 20% absolute gain in cerebral perfusion across the entire ischemic hemisphere, compared to pre-treatment. This result is consistent across the different types of analysis performed and appeared larger (up to approximately 30-35%) in the more densely ischemic regions.

Moreover, our results indicate that infarct growth during the first 24 hours was significantly slowed down by HDT15 application, compared to the flat positioning. This result is also likely to reflect an increased collateral flow induced by HDT15, since the degree of collaterals have been associated with the speed of infarct progression before reperfusion.^30,31^

Previous preclinical studies explored the effect of HDT on stroke outcome.^10–13^ In particular, a pooled analysis on 104 randomized rats from 3 experimental series, including the one from the present study, showed that HDT15 applied 30 minutes after transient MCA occlusion reduced infarct size (relative difference 34%, Cohen’s d 0.56) and improved functional outcome (OR 2.6, 95% confidence intervals 1.1-6.2), compared to flat positioning.^11^

HDT was shown to increase cerebral blood flow using laser doppler^10, 11^ or two-photon microscopy.^13^ We believe that demonstrating an acute beneficial effect of HDT15 on perfusion MRI during MCA occlusion is of utmost importance at a translational level, since perfusion imaging is the current standard for accessing to recanalization therapies.^32^

There is a lack of consensus in current clinical practice regarding the most appropriate head position for patients affected by acute stroke, albeit the most common one is the semi-sitting position at about +30°.^33^ In the large pragmatic trial HeadPoST, flat position had a neutral effect on outcome in acute stroke patients, compared to +30°.^22^ However, recanalization therapies were delivered to a small minority of the HeadPoST participants (approximately 10%). Interestingly, an observational study reported early neurological improvement with early −15° positioning, compared to +30°, in consecutive LVO stroke patients, more than 50% of which received intravenous thrombolysis.^34^

On the contrary, the large clinical trial AVERT showed that early mobilization was associated with neurological worsening, higher mortality and worse neurological outcome in acute ischemic or hemorrhagic stroke.^35^

Interestingly, a recent randomized, pilot clinical trial showed promising results of -20° head down positioning on long-term disability and an excellent safety profile in patients with acute ischemic stroke of predominantly hemodynamic etiology.^36^

Safety of a collateral therapeutics could not be overemphasized. Being aimed to be delivered as an emergency measure to patients with suspected stroke, even in the pre- hospital setting, a collateral therapeutic should be safe not only for acute ischemic stroke, but also in case of intracerebral hemorrhage or other stroke mimics. Available clinical and pre-clinical studies indicated that transient HDT application ranging from 0° to -20° for up to 24 hours was feasible and safe in acute stroke patients, including patients with intracerebral hemorrhage, with no significant additional risk of intracranial hypertension or aspiration pneumonia, and in a rat model of intracerebral hemorrhage.^22,34, 37, 38^

Our study has limitations. First, the sample size of our preclinical trial is small. However, it has been estimated to assess the impact of HDT15 on cerebral perfusion imaging and the primary outcome was met. Second, the baseline collaterals were slightly unbalanced favoring the HDT15 arm and the effect size of the experimental treatment could have been overestimated. However, this is an intrinsic limitation of the unrestricted randomization employed in this study and all between-group comparisons were adjusted for baseline collateral score. Third, results of CBF and CBV have been considered exploratory due to the limits in obtaining AIF reliably, considering the limited spatial resolution of small animal MRI. Fourth, we did not investigate that physiological, biochemical or molecular mechanism of action of HDT15 in experimental LVO stroke, which has still to be defined.

Overall, our results suggest that HDT15 acutely increases cerebral perfusion assessed by perfusion MRI and slowed down infarct growth in experimental LVO stroke. Further research is needed to implement HDT15 as an emergency collateral therapeutic for acute ischemic stroke prior to recanalization therapy.

## ACKNOWLEDGEMENTS

We thank Mr Marco Pini and Mr Jean-Baptiste Langlois for technical assistance.

## AUTHOR CONTRIBUTION STATEMENT

SB, DC, T-HC and FC conceived and designed the study. SB, T-HC, MV, SD, JM, FAP, CP, RB and FC acquired data. SB, DC and EB performed data analysis. SB, CD, T-HC, MW, CF and FB interpreted data. SB, DC, EB and FC drafted the manuscript, which was critically revised and approved by all Authors.

## DECLARATION OF CONFLICTING INTERESTS

The author(s) declared no potential conflicts of interest with respect to the research, authorship, and/or publication of this article.

## FUNDING

The author(s) disclosed receipt of the following financial support for the research, authorship, and/or publication of this article: This work was supported by the Agence Nationale de la Recherche (ANR) [grant number ANR-21-CE17-0028] and the Italian Ministry of University and Research (MIUR) [grant number PRIN 2017CY3J3W].

## SUPPLEMENTARY MATERIAL

Supplemental material for this article is available online.

## Notes

https://openneuro.org/datasets/ds004962/versions/1.0.0

### Competing Interest Statement

The authors have declared no competing interest.

