## Supplementary Material for "Head down tilt 15° increases cerebral perfusion before recanalization in acute ischemic stroke. A pre-clinical MRI study"

Simone Beretta[1,2], Davide Carone[3], Tae-Hee Cho[4,5], Martina Viganò[1], Susanna Diamanti[1], Jacopo Mariani[1], Francesco Andrea Pedrazzini[1], Elisa Bianchi[6], Cristiano Pini[1,7], Radu Bolbos[8], Marlene Wiart[5], Carlo Ferrarese[1,2], Fabien Chauveau[9]

[1]Laboratory of Experimental Stroke Research, Department of Medicine and Surgery, University of Milano-Bicocca, Monza, Italy

[2]Department of Neuroscience, Fondazione IRCCS San Gerardo Monza, Italy

[3]Acute Vascular Imaging Centre, Radcliffe Department of Medicine, University of Oxford, United Kingdom

[4]Department of Vascular Neurology, Hospices Civils de Lyon, Lyon, France

[5]CarMeN Laboratory, INSERM U1060, INRA U1397, INSA-Lyon, Université Claude Bernard Lyon 1, Lyon, France

[6]Istituto Di Ricerche Farmacologiche Mario Negri IRCCS, Milan, Italy

[7]Nuclear Medicine Department, IRCCS San Raffaele Hospital, Milan, Italy

[8]CERMEP-Imagerie du Vivant, Lyon, France

[9]Centre de Recherche en Neurosciences de Lyon, CNRS UMR5292, INSERM U1028, Université Claude Bernard Lyon 1, Lyon, France

### SUPPLEMENTAL MATERIAL

### Supplemental Methods

#### Animal housing and husbandry

Animals were given a minimum of five days to acclimate to the conventional housing facility, under temperature-controlled (range 20–24°C) conditions and a 12 :12 h light-dark cycle, with lights on at 07:00 and off at 19:00. Animals were housed by group of four to six in open polycarbonate cages (Tecniplast, 2000P, L × W × H = 610 × 435 × 215 mm, floor area 2065 cm2), with stainless steel lids. Environmental enrichment included spruce-based bedding of 2–4 mm granulometry (Lignocel 3/4 s), round tinted polycarbonate tunnels (153 × 75 mm, SERLAB), and hazel chew blocks (JR Farm). Animals were given access to pellets of wheat and corn (Teklad Global 18% Protein Rodent Diet, ENVIGO) and tap water ad libitum. During housing, animals were monitored daily for health status.

#### Animal care and monitoring

For pain control, a subcutaneous injection of buprenorphine (a morphine analgesic) is administered 15-20 min prior to surgery. The dose used in rats is 0.01-0.05 mg/kg. Analgesia is supplemented by local application of lidocaine ointment to the surgical field. These treatments can be repeated after awakening and until the end of the experimental protocol, 24 hours after reperfusion, depending on the animal's condition observed. Feeding is facilitated by food (solid or jelly) in the cage.

The project includes an invasive surgical procedure. We therefore consider as humane endpoint any manifestation of second-level pain, and in particular:

- major bleeding (arterial, lasting more than 1 minute) during surgery
- weight loss exceeding 15% during the procedure
- clinical signs of involvement of a territory greater than that of the middle cerebral artery (convulsions, severe hypotonia)
- intracerebral hemorrhage detected on MRI (hyposignals on T2-weighted imaging)
- pain: persistent facial signs (eyes, muzzle, ears, vibrissae) not relieved by buprenorphine treatment (according to Rat Grimace Scale)
- sensorimotor deficits: animal not mobile, not reactive, hypotonic (corresponding to a score <5 on Garcia scale)

The search for these manifestations is carried out when the animal wakes up after the stroke, then at least 2 times during the during the 24-hour protocol (at the end of the day and the following morning).

**Mortality of included animals**

Three rats (all in the flat group) with successful MCA occlusion died several hours (< 24 hours) after completing the perfusion MRI protocol, due to malignant ischemic stroke (verified by necropsy and histology). Data from these animals was used for all available outcome measures.

Five rats (4 in the flat group, 1 in the HDT group) with successful MCA occlusion died < 1 hour after completing the perfusion MRI protocol, during or soon after re-exposure to isoflurane for filament removal. Breath rate was monitored during the MRI protocol and no abnormality was detected in these animals during MRI acquisition. The presumed cause of death was respiratory failure associated with re-exposure to isoflurane after MRI acquisition (SAH and malignant ischemic stroke were excluded by necropsy and histology). Data from these animals were used for the primary outcome only.

**Image processing, region of interest (ROI) and masks**

All image analyses were performed using the Oxford Centre for Functional MRI of the Brain (FMRIB) software library (FSL) unless otherwise specified.

Rigid body registration (6 degrees of freedom) using FMRIB's Linear Image Registration Tool (FLIRT) was used for within time point and across timepoint registration. Brain extraction was performed by applying a manually generated brain mask.

The following mask and ROIs were generated for all included animals:

- Brain mask: two research fellows (JM, MV) manually segmented the brain on the baseline T2 and separately labelled the two hemispheres in an Ischemic hemisphere and Contralateral hemisphere ROIs.
- Infarct core : the ischemic stroke lesion at presentation was automatically segmented on the ADC maps using an externally validated threshold of $530\times10{-}6$ mm2/s;[23] all masks were visually reviewed (SB) and amended if artefacts were present.
- Final infarct: the ischemic stroke lesion was segmented on the 24h T2 by two research fellows (JM, MV).
- Infarct growth was defined by comparing the volume of infarct core with the volume of the final infarct.
- Arterial input function (AIF) mask: the contralateral carotid artery was manually labelled by two manual raters (SD, DC) on the first volume of the DSC-PWI data.

All manual masks were reviewed and any discrepancies were resolved by a third rater.

### Table S1. Technical details of MRI acquisition protocol

| SEQUENCE | Method | TE (ms) | TR (ms) | FOV (cm) | Matrix (pts) | Resolution (µm) | No. Repetitions | B Values (s/mm2) | No. Slices | Slice thickness (mm) | Slice orientation | Acquisition Time |
|---|---|---|---|---|---|---|---|---|---|---|---|---|
| DSC-PWI | EPI | 8.2 | 600 | 3.50 x 3.50 | 80 x 80 | 437 X 437 | 100 | - | 15 | 1 | axial | 1 min |
| ADC-map | EPI-SE | 23.2 | 5000 | 3.50 x 3.50 | 128 x 128 | 273 x 273 | - | 0, 1500, 3000 | 15 | 1 | axial | 4 min 40 s |
| Angio-MRI | Flow Comp | 4.1 | 18 | 3.48 x 2.50 | 256 x 184 | 136 x 136 | - | - | 60 | 0.4 | coronal | 7 m 27 s |
| T2w | SE | 75 | 5000 | 3.50 x 3.50 | 256 x 256 | 137 x 137 | - | - | 15 | 1 | axial | 4 min |

### Table S2. Stroke-related mortality, neuroscore and infarct volume at 24 hours

| | **FLAT** n (%) or mean (SD) | **HDT15** n (%) or mean (SD) | p |
|---|---|---|---|
| Stroke-related mortality | 3/14 (21.4%) | 0/14 (0%) | 0.2222 |
| Garcia neuroscore | 7.1 (5.2) | 10.3 (2.9) | 0.0705 |
| Infarct volume at 24 hours | 341.8 (142.6) | 224.0 (159.7) | 0.0575 |

FLAT = flat positioning. HDT15 = head down tilt 15°. SD = standard deviation.

**Figure S1. Voxelwise analysis of cerebral blood flow (CBF) and cerebral blood volume (CBV)**

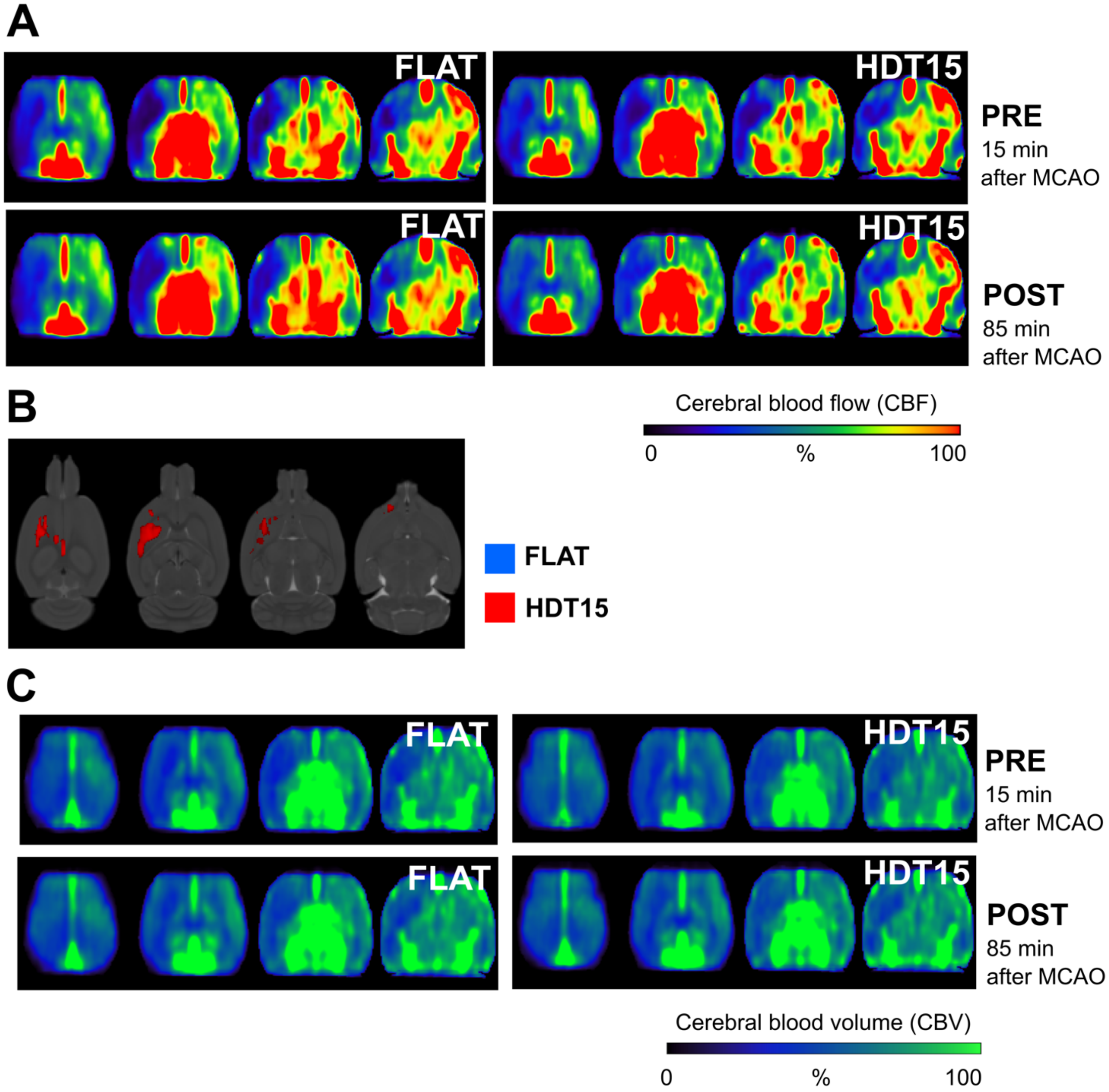
